# Coevolution of a phage cocktail constrains reversible phenotypic resistance in *K. pneumoniae*

**DOI:** 10.64898/2025.12.19.695413

**Authors:** Lucas Mora-Quilis, Pilar Domingo-Calap

## Abstract

The therapeutic potential of phages is frequently compromised by rapid bacterial adaptation. Strategies such as phage cocktails and phage training are widely used to counteract bacterial resistance. Although phage receptor mutations are highly common, bacteria can also exploit flexible non-genetic strategies to evade phage recognition. Overcoming such phenotypic resistance remains particularly challenging due to its transient and reversible nature. Here, we investigated the coevolutionary dynamics between *Klebsiella pneumoniae* and a phage cocktail with complementary host tropisms: a capsule-dependent phage and an acapsular-targeting phage, coevolved with bacteria either individually or in combination. The coevolutionary phage training demonstrated that evolution of the acapsular-targeting phage was the main driver of the enhanced resistance delay observed. Notably, while bacteria could evade individual phages through non-mutational mechanisms, the combined selective pressure imposed by the phage cocktail constrained this reversible escape route, forcing bacteria to evolve mutation-based resistance mechanisms. Together, these findings highlight the importance of elucidating phage-bacteria coevolutionary dynamics to optimize phage-based therapeutics design and reshape bacterial evolutionary trajectories toward favourable outcomes.

**Importance:** Phage therapy is re-emerging as a promising alternative to antibiotics, yet its long-term efficacy is often limited by the rapid evolution of bacterial resistance. While phage cocktails and experimental phage training have been widely proposed to mitigate this problem, most studies focus on genetically fixed resistance, overlooking reversible phenotypic resistance mechanisms. Here, we show that phenotypic resistance based on capsule regulation enables bacteria to evade single-phage treatments without acquiring mutations. In contrast, coevolutionary training of a two-phage cocktail targeting complementary bacterial phenotypes constrains this potentially low-cost escape route, forcing bacteria to adopt a mutation-based resistance, typically associated with higher fitness costs. These findings highlight the importance of phage cocktails not only as therapeutic tools, but also as ecological drivers that redirect bacterial evolutionary trajectories.

## INTRODUCTION

The global rise of antimicrobial resistance has renewed interest in phage therapy as a promising alternative to antibiotics [1–3]. Bacteriophages or phages, bacterial viruses, offer unique advantages such as high specificity, anti-biofilm activity and self-amplification at the site of infection [4, 5]. However, their therapeutic potential is often compromised by the rapid emergence of bacterial resistance [6–8]. Overcoming this evolutionary obstacle requires strategies that identify and improve effective phages to anticipate and constrain bacterial adaptive responses [9–11]. One promising approach is phage training, a directed evolution strategy that exploits the natural adaptability of phages. Classical training protocols typically involve serial passaging on a non-permissive bacterial host to select for mutants with expanded host range [12–14]. In contrast, experimental coevolution provides a dynamic alternative, allowing phages to evolve in parallel with a continuously adapting bacterial population [15–17]. This process mimics natural predator–prey interactions [18, 19], producing phages potentially better equipped to counter diverse resistance mechanisms. Thus, elucidating phage-host interactions and their evolutionary dynamics will provide the basis for the rational design of novel therapeutics and effective bacterial control strategies.

The encapsulated bacterium *Klebsiella pneumoniae* represents an ideal model to study such processes. As a major opportunistic pathogen [20], its polysaccharide capsule is both a key virulence factor and the primary receptor for many phages [21, 22]. Resistance to capsule-dependent phages typically arises through capsule loss [23–25]. Yet, our previous work demonstrated that, this can occur through reversible, non-mutational downregulation of capsule biosynthesis genes, a form of phenotypic resistance that allows transient evasion of phage infection while preserving the ability to restore the capsule once selective pressure ceases [26]. Similar forms of phenotypic resistance have also been described in *Salmonella* spp., *Escherichia coli*, and *Haemophilus influenzae*, suggesting that such reversible strategies may be widespread among clinically relevant pathogens [27–29]. This flexible resistance challenges conventional views of resistance evolution and complicates phage therapy design. Importantly, it also generates exploitable trade-offs: in encapsulated bacteria, capsule loss not only confers resistance to capsule-dependent phages but also exposes alternative receptors such as lipopolysaccharides (LPS) and outer membrane proteins [30–32]. Phage cocktails can exploit these vulnerabilities by targeting multiple receptors simultaneously, a strategy shown to be effective across diverse bacterial systems [33, 34]. Dual-receptor targeting constrains the bacterial population, substantially delaying the emergence of resistance in the short term [23, 35, 36]. However, the long-term efficacy of phages against reversible phenotypic resistance, and the resulting phage-bacteria coevolutionary dynamics, remains poorly understood.

To address this, we coevolved a phage cocktail composed of the capsule-dependent phage Cap62 and the acapsular-targeting phage CuaHET1 [26, 35] with the reference capsular type-1 *K. pneumoniae* strain across 20 passages. This experiment allowed us to evaluate the ability of evolved phages to delay resistance, and to characterize the genetic and phenotypic bases of bacterial adaptation under combined phage pressures. Our results showed that coevolutionary training enhances cocktail efficacy and drives *K. pneumoniae* toward new, genetically fixed resistance mechanisms involving modifications of the LPS core, revealing the multilayered nature of host–phage evolution.

## RESULTS

### Bacteria and phages coevolve and persist over 20 passages

To explore how coevolution can both optimize phage efficacy and shape bacterial evolutionary trajectories, we coevolved the phages Cap62 and CuaHET1 with the reference capsular type-1 *K. pneumoniae* in mono- and co-inoculation (cocktail) across three independent lineages for 20 passages (Figure 1A). Both bacterial and phage populations persisted throughout the experiment. Population densities were measured during the first 10 passages to monitor dynamics. Bacterial densities ranged from 3.67 × 10^7^ colony forming units (CFU)/mL to 9.00 × 10^9^ CFU/mL (Figure 1B). Notably, bacteria coevolved in the presence of both phages remained at significantly lower densities with an average of 7.70 × 10^8^ CFU/mL for the three linages (Kruskal-Wallis, *p* < 0.001; Dunn’s test, *p-adj* < 0.001). Phage densities varied between samples. In general, phage Cap62 persisted at high frequencies in both mono- and co-inoculations, ranging from 2.80 × 10^7^ to 2.17 × 10^10^ plaque forming units (PFU)/mL, with a mean of 2.46 × 10^9^ and 2.10 × 10^9^ PFU/mL, respectively (Figure 1C). However, phage CuaHET1, although persistent throughout the evolutionary process, had a significantly lower titer in the mono-inoculation condition compared to the cocktail condition (1.92 × 10^7^ and 1.73 × 10^9^ PFU/mL, respectively; T-test, *p* < 0.001).

**Figure 1.**
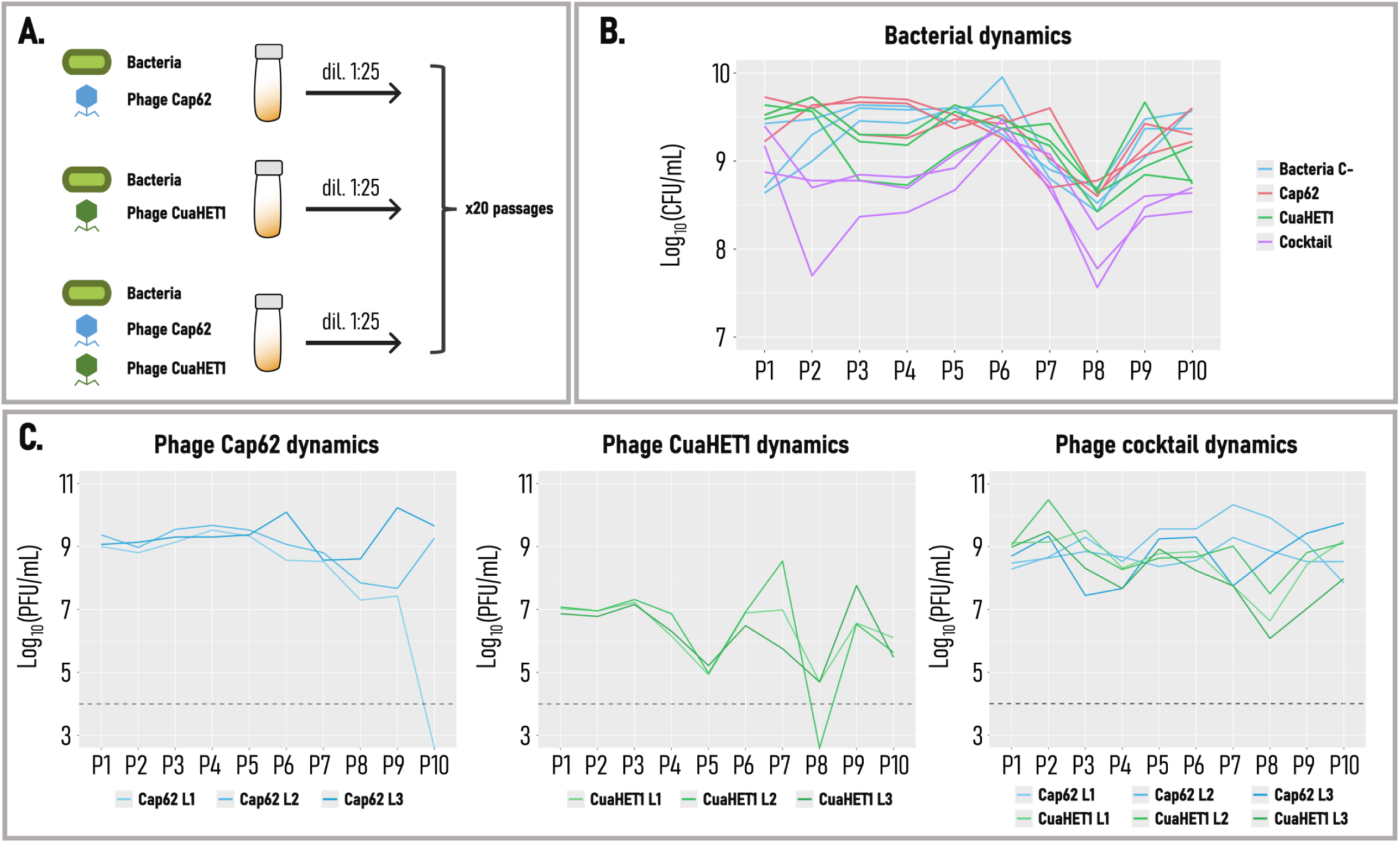
Bacteria-phage co-evolution system. **A.** Schematic representation of the co-evolution experiment. Bacteria were infected with phage Cap62 and CuaHET1, either in mono- or co-inoculation (cocktail), and propagated for 20 passages at a 1:25 dilution in fresh medium. **B.** Bacterial dynamics during the first 10 passages of the co-evolution, expressed as CFU/mL. Data from three independent lineages are shown for each condition. **C.** Phage dynamics during the first 10 passages, expressed as PFU/mL. Data from three independent lineages are shown. The grey dotted line represents the detection limit.

### Evolved phages exhibit improved activity in delaying resistance

To determine whether phage evolution enhanced their capacity to delay the emergence of bacterial resistance, supernatants from the 20th passage of the coevolution experiment were used to infect the ancestral bacterial culture and compared with the ancestral phages (Figure S1). Growth curves revealed that all evolved phages tended to delay resistance relative to their ancestral counterparts, with the most pronounced effect observed for phages evolved under the cocktail condition. To investigate this effect in greater detail, five independent plaques of Cap62 and five of CuaHET1 evolved in the cocktail condition were randomly isolated from each of the three evolutionary lineages. These evolved isolates were then combined in Cap62–CuaHET1 pairs, either mixing evolved phages with each other or with their complementary ancestral variant. These combinations were used to infect the ancestral bacterial culture, and their ability to delay resistance was quantified using the resistance delay index (RDI). The evolved phages exhibited varying degrees of improvement across the three independent coevolution lineages (Figure 2A). In lineage 1, the RDI ranged from 0.485 to 0.858, compared to the ancestral combination (Cap62_Anc_ + CuaHET1_Anc_) which showed a value of 0.616. Similarly, in lineage 3, RDI ranged from 0.406 to 0.918, while the ancestral cocktail reached only 0.416. In both cases, all combinations involving evolved Cap62 and evolved CuaHET1 yielded higher indices than the ancestral pair, indicating that coevolution enhanced the ability of the cocktail to delay resistance. In contrast, lineage 2 showed limited improvement, with values ranging from –0.643 to 0.685, compared to the ancestral combination of 0.597.

**Figure 2.**
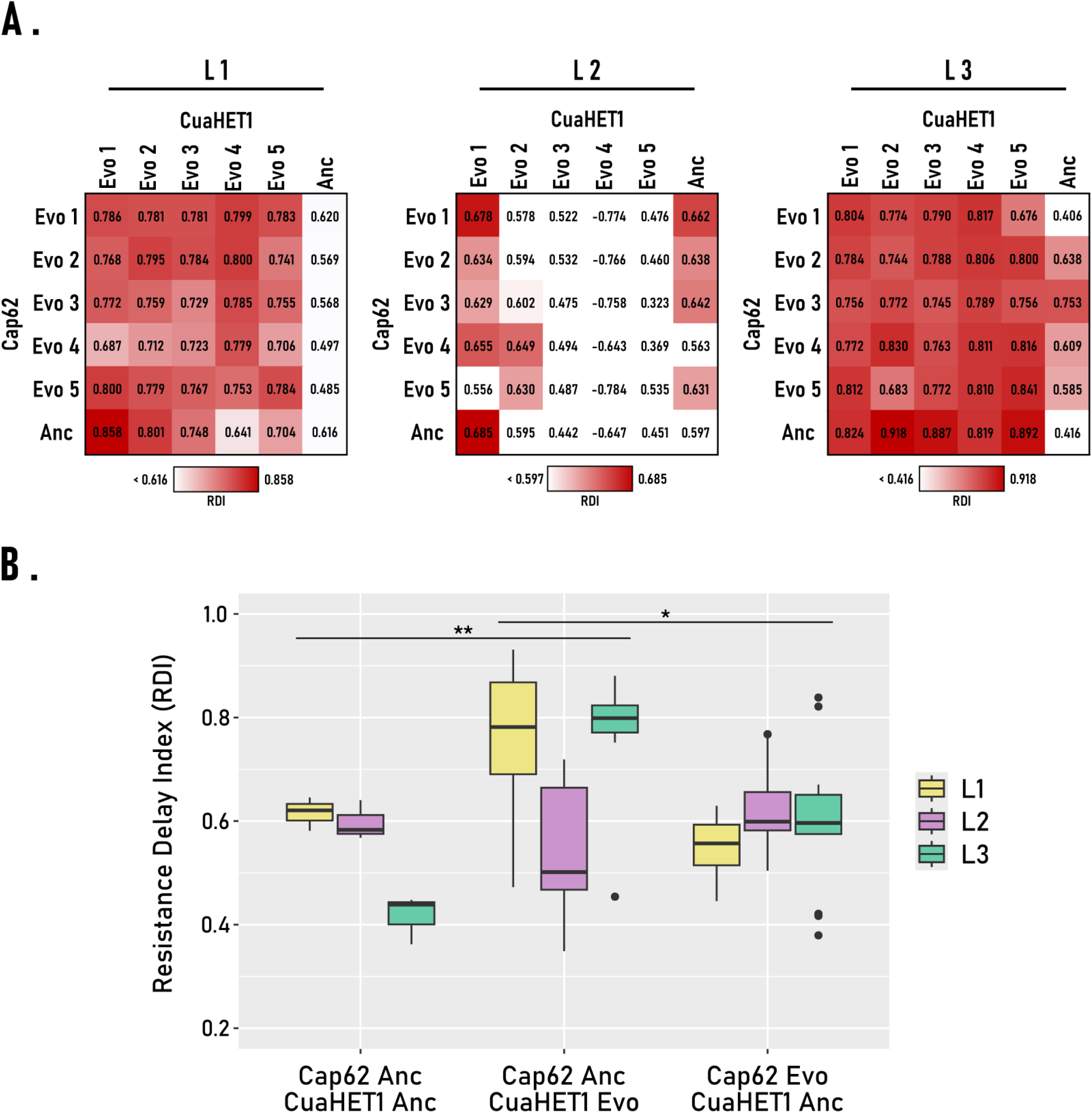
Ability of evolved phages Cap62 and CuaHET1 from the cocktail condition to delay resistance. **A.** Combinations of evolved Cap62 and evolved CuaHET1 phages from the cocktail condition across the three evolutionary lineages, showing the resistance delay index (RDI) for each combination. Evo denotes evolved phage isolates, whereas Anc refers to the ancestral phage variants. The colour scale represents the RDI value, where white corresponds to the minimum value, defined as the combination of ancestral Cap62 and ancestral CuaHET1 phages, and red represents the maximum RDI. Values equal to or below the minimum are displayed in white. **B.** RDI values grouped by specific phage combinations for each lineage. The combination of ancestral Cap62 with evolved CuaHET1 displayed significantly higher RDI values than either the ancestral pair or the combination of evolved Cap62 with ancestral CuaHET1 phage (Kruskal-Wallis test followed by Dunn’s post hoc test, *p-adj* < 0.05 (*), *p-adj* < 0.01 (**)). For clarity, six RDI values corresponding to the combination Cap62_Anc_ + CuaHET1_Evo4_ (L2) are not shown but were included in the statistical analysis.

### The evolution of phage CuaHET1 is primarily responsible for the improved delay on resistance

To determine the contribution of each phage to the enhanced resistance delay, we compared the performance of ancestral-evolved combinations. The highest RDI values were observed for combinations of ancestral Cap62 with evolved CuaHET1, which were significantly higher than those of the ancestral pair (Kruskal-Wallis, *p <* 0.01; Dunn’s test, *p-adj <* 0.05) (Figure 2B). In contrast, combinations of evolved Cap62 with ancestral CuaHET1 yielded RDI values similar to the ancestral combination. These findings indicated that CuaHET1 evolution contributed more strongly to the enhanced performance of the phage cocktail. To better illustrate the impact of evolved phages on bacterial dynamics, we plotted representative growth curves from the best-performing combinations of the three lineages (Figure 3). These curves visually confirmed the patterns observed with the resistance delay index. Whereas phages from lineage 2 showed no improvement in delaying resistance compared to the ancestral cocktail, combinations from lineages 1 and 3 that included evolved CuaHET1 achieved a prolonged suppression of bacterial growth. The strongest effect was observed when evolved CuaHET1 was combined with the ancestral Cap62, almost completely preventing bacterial growth. Conversely, combinations containing evolved Cap62 with ancestral CuaHET1 exhibited the weakest performance, with resistance emerging earlier than in cultures treated with the ancestral cocktail.

**Figure 3.**
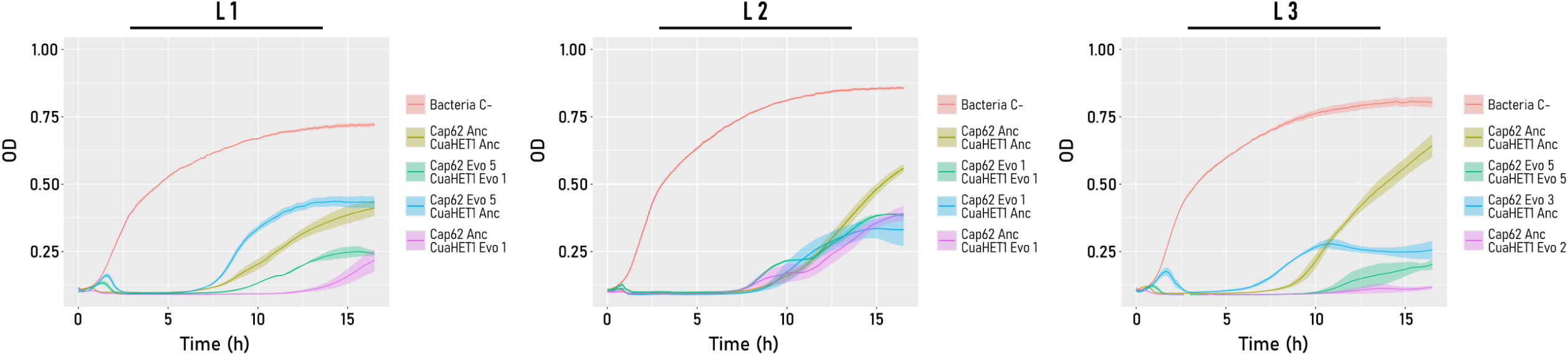
Infection dynamics of the most effective evolved Cap62 and CuaHET1 phage combinations from the cocktail condition. Bacterial growth curves are shown for the control (Bacteria C-) and for infections with the Cap62 and CuaHET1 combinations displaying the highest resistance delay index (RDI), including combinations of evolved (Evo) and ancestral (Anc) phages. Shaded areas represent the standard error of the mean (SEM).

### Phages evolve mainly via mutations in receptor binding proteins

To identify the genetic basis of phage adaptation during coevolution, we analyzed the genomic changes accumulated after 20 passages of evolution under mono-inoculation and cocktail (co-inoculation) conditions (Data S1). All samples were successfully sequenced, except for phage Cap62 in lineage 2 of the cocktail. Phage Cap62 accumulated four identical mutations in both mono-and co-inoculation in the tail fiber protein (CDS 0007) (alternative allele frequency > 80%) (Figure 4A), as well as an additional mutation in a hypothetical protein (CDS 0022), with unknown function according to BLASTp and a LIM-like domain according to HHPred. Phage CuaHET1 accumulated mutations mainly in structural and recognition genes such as the hoc-like head decoration (CDS 0016), the long tail fiber protein proximal connector (CDS 0085), tail fiber adhesin (CDS 0088), the baseplate wedge subunit (CDS 0280) and the hypothetical protein (CDS 0229) (freq. > 70 % in at least two linages). The only gene mutated either in mono- and co-inoculation in the three linages was the tail fiber protein (CDS 0087) which showed a C → T substitution in the position 60709 (mono-inoculation) (alternative allele frequency = 1), and a G → T substitution in the position 61128 (co-inoculation) (alternative allele frequency = 100%) (Figure 4B). The tail fiber adhesin (CDS 0088) also was mutated in both mono- and co-inoculation with at least two linages with an alternative allele frequency > 92%.

**Figure 4.**
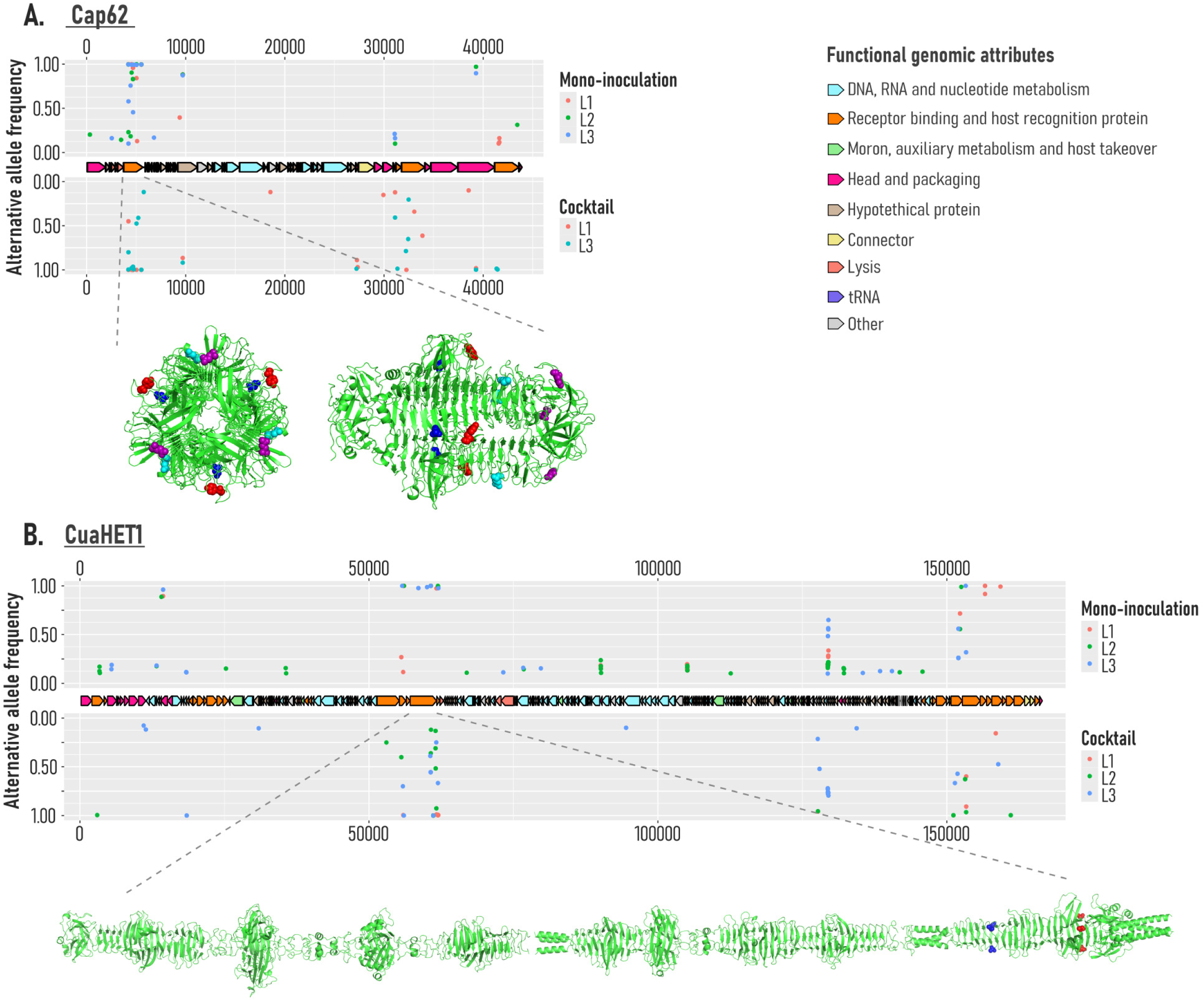
Genomic and structural changes in evolved phages Cap62 and CuaHET1. Alternative allele frequencies (≥ 10%) across the genomes of phages Cap62 (A) and CuaHET1 (B) for the three independent lineages evolved under mono-inoculation and cocktail conditions. Genomic functional categories are represented by coloured boxes. Below, structural representations of the Cap62 tail fiber protein (CDS 0007) are shown in transversal (left) and longitudinal (right) views, highlighting amino acids changes that were identically detected under both mono-inoculation and cocktail conditions. For CuaHET1, the longitudinal structure of the tail fiber protein (CDS 0087) is displayed, indicating the mutations detected in both conditions.

### Bacteria evolve via diverse mechanisms to mono- and co-inoculation

To characterize bacterial adaptation to different phage pressures, we examined phenotypic and genomic changes after 20 passages of coevolution. Bacteria coexisted with phages throughout the experiment, although their phenotypes differed depending on the infection condition. Bacteria evolving in absence of phages and in CuaHET1 mono-inoculation retained the capsular phenotype, with the latter increasing the mucoid appearance (Figure 5A). In contrast, bacteria evolving in the presence of Cap62, either in mono- and co-inoculation, developed an acapsular phenotype and exhibited a reduced mucoid appearance. Whole-genome sequencing revealed that bacteria coevolved with either Cap62 and CuaHET1 in mono-inoculation displayed no detectable mutations, whereas only those coevolved with the phage cocktail accumulated mutations in two independent lineages (Figure 5B). In lineage 2, six genes with diverse functions were mutated, including those related to efflux-mediated drug resistance (*emrA*), carbon metabolism regulation (*iclR*), cell wall remodelling (*amiA*), and tRNA biogenesis. Notably, a mutation was also identified in *waaE* (CDS 0136), a glucosyltransferase involved in the biosynthesis of the inner core of the LPS. In lineage 3, a mutation occurred in a hypothetical protein (CDS 0145) predicted possess sialyltransferase activity and located immediately adjacent to *rfaC* (CDS 0146) and *rfaF* (CDS 0147) within the LPS biosynthetic cluster. The proximity and functional context of these mutations strongly suggest that the phage cocktail exerts selective pressure on LPS structure, promoting the emergence of LPS-associated resistance variants.

**Figure 5.**
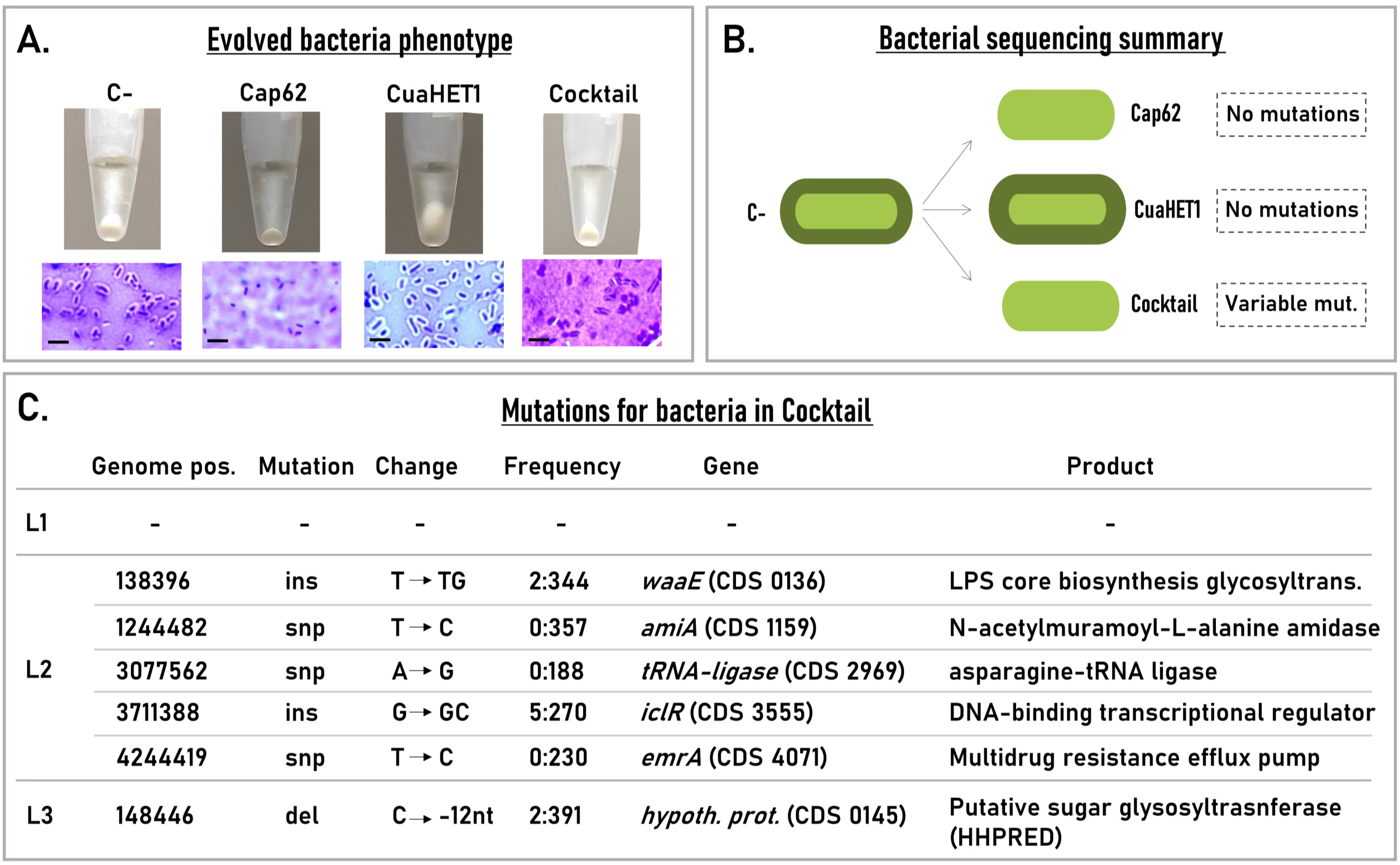
Evolved bacteria. **A.** Evolved bacterial phenotypes. Mucoidy of bacterial culture pellets after centrifugation for bacteria evolved in LB (C-), mono-inoculation (Cap62 or CuaHET1), and co-inoculation (cocktail). Microscopy images show the presence of the capsule in C- and Cua conditions. Scale bars represent 5 µm. **B.** Summary of bacterial genome sequencing. Mutations were detected only in the cocktail condition. **C.** Mutations in bacteria evolved with the phage cocktail. The lineage (L1, L2, L3), the genome position (bp) and mutation type are indicated (ins: insertion, snp: single-nucleotide polymorfism, del: deletion). Nucleotide changes and their respective frequencies compared to the consensus allelle are shown. Genes, corresponding CDS based on genome annotation, and the names of the protein products are listed.

## DISCUSSION

Coevolutionary experiments offer an interesting framework to study the interplay between phages and bacteria, which is highly relevant for the design of therapeutic cocktails. In our study, the coevolution of a phage cocktail against *K. pneumoniae* not only enhanced its efficacy but also redirected the bacterial evolutionary trajectory. The contrast between single-phage and cocktail coevolution reveals a hierarchical organization of bacterial defenses governed by evolutionary costs [37, 38]. When exposed to individual phages, the phenotypic regulation of the capsule may act as a first, low-cost, and reversible line of resistance [28, 39]. By transiently modulating capsule expression, bacteria can evade infection without permanently compromising essential cell envelope structures. While capsule down-regulation prevents recognition by capsule-targeting phages [31, 32], up-regulation can shield alternative surface receptors such as LPS or OMPs, hindering adsorption by non-capsule-targeting phages. In contrast, simultaneous exposure to both phages made this reversible mechanism insufficient, triggering the emergence of mutations in essential LPS biosynthetic genes such as *waaE* [40]. A similar pattern has been reported in *Salmonella* spp., where phenotypic phage resistance persisted until the coevolving phage acquired the ability to infect both ancestral and resistant hosts, ultimately driving the emergence of genetically fixed resistance [41]. Our results suggest that LPS may act as receptor for CuaHET1, and its mutation provide an alternative escape route when capsule modulation alone fails. However, these mutations are likely associated with substantial fitness costs, as they can compromise outer membrane integrity and altering interactions with the host environment [40, 42]. Therefore, only when the phenotypic route is exhausted does the bacterium resort to genetically fixed resistance.

Phage adaptation to bacterial hosts is frequently mediated by mutations in the receptor-binding proteins (RBPs), a common mechanism for modulating host range and infection efficiency [13, 14, 43]. For instance, the *Pseudomonas aeruginosa* phage PWJ acquired a SNP in the tail fiber protein that enabled it to suppress phage resistance in a murine long infection model [44]. Similarly, previous work showed that evolution of phage U136B led to increased adsorption to *E. coli* through mutations in the tail fiber protein [45]. The selective pressures driving RBP evolution depend strongly on the bacterial phenotype encountered. For CuaHET1, mutations differed between mono- and co-inoculation, reflecting adaptation to capsulated and acapsular hosts, respectively. These changes likely affected RBPs involved in LPS recognition, imposing a selective pressure on bacterial LPS biosynthetic genes, particularly in the cocktail condition, where the loss of capsule exposes the LPS. In contrast, Cap62 accumulated identical RBP mutations in both scenarios, consistent with adaptation to similar acapsular hosts. Thus, while the cocktail was designed to exploit complementary tropisms, phage evolution converged toward adaptation to infect acapsular variants. Previous work with phage λ demonstrated that destabilizing mutations can relax RBP structure enabling recognition of new receptors [46, 47]. Although Cap62 does not appear to exhibit such receptor switching, its RBP mutations may instead reduce depolymerase activity to cope with lower or altered capsule, but at the cost of decreased infectivity toward the ancestral capsulated strain.

While from a functional perspective, the adaptive changes in Cap62 appear to be ineffective, from an ecological standpoint, the role of Cap62 remained essential. Cap62 acted as a driver selecting the emergence of acapsular variants, supplying continuously susceptible hosts for CuaHET1, which exploited this newly dominant phenotype. In this sense, Cap62 shaped the ecological landscape, while CuaHET1 adapted to it. Previous work has shown that phage fitness landscapes are continuously reshaped by reciprocal bacterial adaptation [48], and that increased phage genotypic diversity can constrain bacterial arms-race evolution compared to single-phage coevolution [49]. Consistently, the simultaneous presence of Cap62 and CuaHET1 limited the ability of *K. pneumoniae* to rely on reversible capsule modulation, forcing more permanent and costly resistance mechanisms. This reciprocal interplay illustrates the complex ecological interactions that can emerge within multi-phage systems, where one phage may indirectly facilitate or hinder the success of another phage by reshaping the bacterial population structure [50, 51]. Therefore, understanding these interactions, particularly under reversible forms of resistance, is essential to elucidate the long-term evolutionary dynamics and ecological stability of phage-bacteria systems. It would be interesting to investigate whether such ecological dynamics occur not only under laboratory conditions but also in vivo. Pathogens with high phenotypic plasticity, such as *K. pneumoniae* [52, 53], may hinder single-phage treatments through reversible modulation of capsule expression. In such cases, phage cocktails targeting complementary bacterial phenotypes, including reversible resistant variants, can block this flexible escape route, forcing the bacterium toward costlier and permanent adaptations. These findings support the advantage of phage cocktails over monotherapy or sequential treatments, in these particular situations.

Future work should explore these dynamics in structured or host-associated environments, and at the molecular level, define how specific receptor-binding protein changes and LPS modifications determine infection outcomes. By integrating these evolutionary insights, phage therapy can be guided not only to treat infections effectively, but also to shape bacterial evolution in therapeutically favorable directions.

## METHODS

### Phages and bacterial strains

The bacterial strain used in this experiment is the *K. pneumoniae* NCTC5054 (CABDWB010000000), the reference strain for capsular serotype K1 from the Statens Serum Institute (Copenhagen, Denmark). The strain was previously characterized [26], and its raw sequencing data have been deposited in the European Nucleotide Archive (ENA) under accession number PRJEB97490. The phages used in this study are phage vB_Kpn_Cap62 (Cap62) (PQ741857) and vB_Kpn_CuaHET1 (CuaHET1) (PX060900), which were previously characterized [26, 35]. *K. pneumoniae* serotype K1 NCTC5054 was used as the propagation host for Cap62, whereas *K. pneumoniae* serotype K60 (NCTC9180) was used for phage CuaHET1. Phage and bacterial cultures were routinely propagated in lysogeny broth (LB) supplemented with 3.78 mM CaCl_2_ at 37°C with shaking.

### Phage-bacteria coevolution

Coevolution experiments were performed by propagating *K. pneumoniae* serotype K1 with phage under different infection conditions: bacteria infected with phage Cap62 at a multiplicity of infection (MOI) of 0.1; bacteria infected with phage CuaHET1 at an MOI of 0.1; bacteria infected with both Cap62 and CuaHET1 (cocktail) at a combined MOI of 0.2; and bacteria without phages as a control in the absence of phage. Each condition was propagated in triplicate, yielding three independent evolutionary lineages per treatment. Cultures were maintained in a final volume of 2.5 mL, and at each transfer, 100 μL of the previous culture were inoculated into fresh medium. A total of 20 serial passages were performed. Phage and bacterial densities were determined during the first 10 passages by titration of phage particles and bacterial cells, respectively. From every passage, aliquots were stored at –70°C for subsequent phage and bacterial genome sequencing.

### Isolation and purification of evolved phages

Single phage isolates were obtained from the 20th passage of the cocktail condition for each evolutionary lineage. Cultures were centrifuged (5 min, 8000 × g, 4°C), and the supernatant was plated on *K. pneumoniae* serotype K1 for Cap62 isolation or on serotype K60 for CuaHET1 isolation. After incubation for 16 h at 37°C, five individual lysis plaques of evolved Cap62 and five of evolved CuaHET1 were randomly selected. Each plaque was resuspended in 4 µL of LB supplemented with CaCl₂ and mixed thoroughly using a pipette. To purify each isolate, tenfold serial dilutions were prepared and plated onto their respective host strains, and the plaque isolation procedure was repeated a total of three consecutive times. The resulting purified evolved phages were then amplified in their corresponding hosts and stored at –70°C.

### Infection growth curves and resistance delay index

To evaluate the ability of phages to delay the emergence of bacterial resistance, infections were conducted in liquid culture and monitored through optical density (OD) measurements. Combinations were made between individual isolates of evolved Cap62 and evolved CuaHET1, as well as combinations including the ancestral variants. Experiments were performed in 96-well plates with a final volume of 150 µL per well. Each well contained approximately 10⁷ plaque-forming units (PFU)/mL. When the two phages were combined, each was added at half the concentration (5 × 10⁶ PFU/mL; multiplicity of infection (MOI) = 0.5 per phage). Cultures were incubated for 16 h at 37°C with shaking, and optical density at 620 nm was automatically recorded using a Multiskan FC plate reader. To quantitatively compare the ability of each phage or phage combination to delay the emergence of bacterial resistance, we calculated a Resistance Delay Index (RDI) using the following formula: RDI = 1 - (OD_final_ / OD_initial_ × T_bmax_)), where OD_final_represents the optical density at the end of the experiment, OD_initial_ is the initial optical density of the culture, and T_bmax_corresponds to the time (h) at which the culture reaches the maximum slope in the OD curve. Consequently, higher RDI values indicate stronger suppression of bacterial growth.

### Capsule staining and microscopy

Capsule presence in bacterial cultures was assessed by contrast staining with crystal violet and nigrosine. Briefly, liquid cultures with a minimum concentration of 5 × 10⁹ CFU/mL were fixed for 20 min with 2.5% formaldehyde in the presence of 100 mM lysine. After fixation, cells were pelleted by centrifugation, washed with 1× phosphate-buffered saline (PBS), and 25–40 µL of the suspension was mixed with a drop of 10% nigrosine on a glass slide. The mixture was smeared along the slide and air-dried. The preparation was then covered with 1% crystal violet for 5 min at room temperature, gently rinsed with distilled water, air-dried, and examined under a light microscope.

### Genomic characterization of evolved phages and bacteria

To characterize the genetic changes occurring during coevolution, both the phage and bacterial fractions from passage 20 of each evolutionary condition and lineage were processed for sequencing. Cultures were centrifuged at 8000 ×g for 10 min at 4°C, and the supernatant and pellet were used for phage and bacterial DNA extraction, respectively. For phage sequencing, supernatants were filtered through 0.22 µm and concentrated using the Concentrating Pipette Select (Innovaprep). Concentrated samples were treated with DNAase, proteinase K (20 mg/mL) and SDS (10%) to release viral nucleic acids from capsids, which were then purified using the Maxwell RSC Instrument (Promega). For bacterial sequencing, pellets were washed twice with 1× PBS, and total genomic DNA was extracted using the PureLink Genomic DNA Mini Kit (Invitrogen). Phage and bacterial DNA libraries were prepared with the Nextera XT DNA Kit (2 × 150 bp), and sequenced using the Illumina NextSeq 550 platform. Read quality was assessed with FastQC [54]. Single-nucleotide variants (SNVs) in the evolved phages were identified using LoFreq [55] with the ancestral phage genomes as reference. Alternative allele frequencies were represented in R using ggplot2 [56] and gggenomes [57] packages. SNVs in the bacterial populations were predicted by Snippy [58], and the potential functions of hypothetical proteins were inferred using HHpred [59].

### Structural modeling and visualization of tail fiber proteins

The three-dimensional structures of the tail fiber proteins of phages Cap62 (CDS 0007) and CuaHET1 (CDS 0087) were predicted using AlphaFold 3 [60] via the Protein Structure Prediction Server (https://alphafoldserver.com/). Both proteins were predicted as trimeric complexes. The resulting PDB files were visualized and analyzed using PyMOL v3.0.5 (Schrödinger, LLC). The ancestral protein structures were represented, and the amino acid positions corresponding to the mutations detected in the evolved variants were highlighted as dots to facilitate structural interpretation.

## AUTHOR CONTRIBUTIONS

Conceptualization, L.M-Q. and P.D-C.; investigation, L.M-Q.; supervision, P.D-C., writing—original draft, L.M-Q.; writing—review & editing P.D-C.; funding acquisition, P.D-C.

## ACKNOWLEDGMENTS

We would like to thank Tamara Barcos for technical assistance. This research was funded by project PID2020-112835RA-I00 and PID2023-150309OB-I00 funded by MCIN/AEI /10.13039/501100011033, and project SEJIGENT/2021/014 funded by Conselleria d’Innovació, Universitats, Ciència i Societat Digital (Generalitat Valenciana) to P.D-C. P.D-C. was financially supported by a Ramón y Cajal contract RYC2019-028015-I funded by MCIN/AEI/10.13039/501100011033, ESF Invest in your future. L.M-Q. was funded by a PhD fellowship FPU19/04611 from Spanish MCIU.

## DATA AVAILABILITY

The genome sequences of phage vB_Kpn_Cap62 (Cap62) and vB_Kpn_CuaHET1 (CuaHET1) are publicly available in GenBank under accession numbers PQ741857 and PX060900, respectively. Raw sequencing data for the evolved phages and bacterial populations will soon be publicly available in the European Nucleotide Archive (ENA).

---

**Figure S1.**
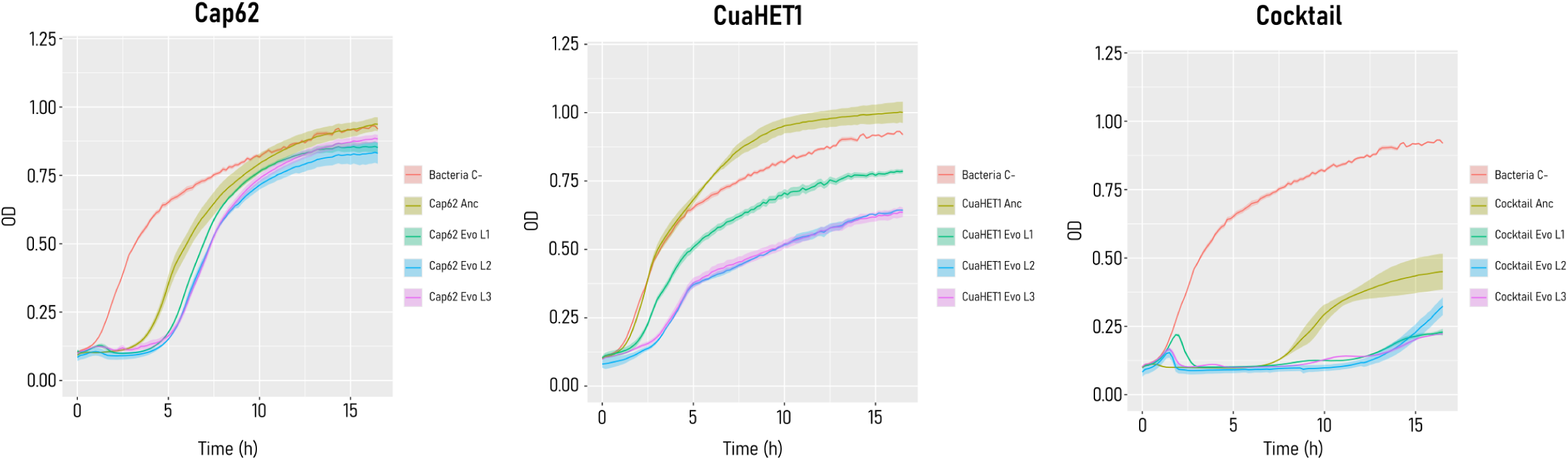
Evolved phage population infection dynamics. Bacterial growth curves of the ancestral strain exposed to the evolved phage populations (lineages L1-L3) from the 20^th^ passage of each evolution condition (Cap62, CuaHET1 and cocktail), compared with their respective ancestral phages. Shaded areas represent the standard error of the mean (SEM).

**Data S1. Phage mutations.** Mutations with an alternative allele frequency ≥ 10% for Cap62 mono-infection, Cap62 cocktail, CuaHET1 mono-infection, and CuaHET1 cocktail. Each condition is presented in a separate worksheet of the .xlsx file.

